# Plasma Activated Water as a Pre-Treatment Strategy in the Context of Biofilm-Infected Chronic Wounds

**DOI:** 10.1101/2023.07.26.550769

**Authors:** Heema K. N. Vyas, Binbin Xia, David Alam, Nicholas P. Gracie, Joanna G. Rothwell, Scott A. Rice, Dee Carter, Patrick J. Cullen, Anne Mai-Prochnow

## Abstract

Healing and treatment of chronic wounds are often complicated due to biofilm formation by pathogens. Here, the efficacy of Plasma Activated Water (PAW) as a pre-treatment strategy has been investigated prior to the application of topical antiseptics polyhexamethylene biguanide, povidone iodine, and MediHoney, which are routinely used to treat chronic wounds. The efficacy of this treatment strategy was determined against biofilms of *Escherichia coli* formed on a plastic substratum and on a human keratinocyte monolayer substratum used as an *in vitro* biofilm-skin epithelial cell model. PAW pre-treatment greatly increased the killing efficacy of all the three antiseptics to eradicate the *E. coli* biofilms formed on the plastic and keratinocyte substrates. However, the efficacy of the combined PAW-antiseptic treatment and single treatments using PAW or antiseptic alone was lower for biofilms formed in the *in vitro* biofilm-skin epithelial cell model compared to the plastic substratum. Scavenging assays demonstrated that reactive species present within the PAW were largely responsible for its anti-biofilm activity. PAW treatment resulted in significant intracellular RONS accumulation within the *E. coli* biofilms, while also rapidly acting on the microbial membrane leading to outer membrane permeabilisation and depolarisation. Together, these factors contribute to significant cell death, potentiating the antibacterial effect of the assessed antiseptics.

## 1. Introduction

In Australia, non-healing chronic wounds (burns, pressure ulcers, diabetic foot ulcers, venous leg ulcers etc) annually costs the healthcare system $3.5 billion; in the United Kingdom, chronic wound care costs £5.3 billion per year; and in the United States, this figure alarmingly exceeds $28 billion [1, 2]. Various pathogens colonise and contaminate chronic wounds such as Gram-positive (*Staphylococcus aureus, Enterococcus faecalis, Streptococcus agalactiae*) and Gram-negative (*Escherichia coli* and *Pseudomonas aeruginosa*) bacteria and fungi (*Candida albicans* and *Aspergillus fumigatus*) [3, 4]. Each of these pathogens are prolific biofilm formers, which can delay healing, complicate treatment, and contribute to the recalcitrant and recurrent nature of chronic wounds [4]. Biofilms are microbial assemblages that can aggregate on a surface and are typically found embedded within a self-produced and/or host-derived protective matrix of extracellular polymeric substances (EPS) [4]. Biofilms are difficult to clear via host immunity and display increased antimicrobial tolerance, and many currently available antimicrobials do not specifically target biofilms [4].

Worryingly, several antimicrobials have been deemed redundant as their overuse and overreliance has resulted in the rapid increase and emergence of antimicrobial resistance (AMR). In wound care, this has seen a shift from topical and systemic antibiotic use towards topical antiseptic ointments, creams, foams, and wound rinses/soaks. Topical antiseptics like polyhexamethylene biguanide (PHMB), povidone iodine (PI), and medical-grade honey are widely recognised first-line treatments, that non-selectively reduce, inhibit, or eradicate microorganisms associated with critically colonised wounds. Despite their promise as safe, cheap, easily appliable, broad-spectrum antimicrobial agents, evidence of their anti-biofilm activity is limited [5].

Plasma medicine is a science that has been investigated in the biomedical field and in clinical practice for cosmetic purposes, cancer therapy, and the treatment of various infections (fungal nails, dental plaque, infected root canals etc,.) [6, 7]. Plasma medicine involves the application of cold atmospheric plasma (CAP) for therapeutic purposes, either directly to the wound or by generating plasma activated liquids [8]. The highly reactive environment of CAP contains several charged particles (electrons, negative and positive ions), excited atoms and molecules, radical species, and UV-photons, which have antimicrobial activity [8]. Interfacing CAP directly with water can transfer these reactive species, generating plasma activated water (PAW). PAW has demonstrated potent antimicrobial activity thought to arise from the variety of short- and long-lived reactive oxygen and nitrogen species (RONS) [9]. PAW is an effective alternative to traditional antimicrobials, and as it acts on multiple targets opportunities for resistance are reduced [10]. PAW has demonstrated antimicrobial efficacy against various planktonic Gram-positive and Gram-negative bacteria, fungi, and viruses [11, 12]. However, its anti-biofilm efficacy remains underexplored.

Here, we have assessed the efficacy of PAW as a pre-treatment strategy to improve the anti-biofilm activity of routinely used topical chronic wound antiseptics PHMB, PI, and medical-grade honey. To aid the translation from the lab to clinical use, we have assessed the anti-biofilm activity of this strategy in an *in vitro* biofilm model that includes a keratinocyte monolayer to mimic the substratum of the wound bed. Inclusion of the host cells is important because biofilms that are formed in simple *in vitro* model systems (e.g., reliant upon plastic, glass, or steel surfaces) lack the impact of host factors, subsequently affecting biofilm antimicrobial susceptibility profiles [13]. We demonstrate that PAW initially kills a significant portion of biofilm cells, and subsequent application of antiseptics results in complete biofilm eradication. Lastly, the mechanisms underpinning PAWs anti-biofilm activity were also investigated. Overall, our findings support further investigation into PAW as a component in wound care, with PAW pre-treatment potentiating the anti-biofilm activity of routinely used topical antiseptics.

## 2. Materials and Methods

### 2.1. Strain and Culture Conditions

*Escherichia coli* has been identified as a common biofilm former in chronic wounds and has thus been selected for this study [3]. *E. coli* (ATCC 25922) was routinely maintained on Luria-Bertani (LB) agar (10.0 g/L tryptone (pancreatic digest of casein), 5.0 g/L yeast extract powder, 10.0 g/L sodium chloride, and 7.5 g/L agar) and cultured in liquid LB media at 37°C at 160 rpm.

### 2.2. Human Keratinocyte Cell Culture Conditions and Monolayer Formation for the *In Vitro* Biofilm-Skin Epithelial Cell Model

HaCaTs, a human epidermal keratinocyte cell line (CLS Cat# 300493/p800_HaCaT), was cultured and maintained in Dulbecco’s Modified Eagle Medium (DMEM) F12 (Gibco, USA), supplemented with 2 mM L-glutamine (Gibco, Life Technologies, USA) and 10% (v/v) heat-inactivated foetal bovine serum (Bovogen Biologicals, Australia) at 37°C in 5% CO_2_ and 20%O_2_ atmospheric conditions. HaCaT keratinocyte cell monolayers were generated as per modified methods of Vyas [14] to encompass host factor presence in the *in vitro* biofilm-skin epithelial cell model. In brief, wells of 96-well flat bottom microtiter plates were pre-coated with 300 µg/mL collagen I from rat tail (Gibco, Life Technologies, USA) for 1 h at 37°C. Once coated, excess collagen was removed, and wells were washed with sterile 1×PBS. Then, each well was seeded with 150 µL HaCaT cell suspension (≈1×10^6^ cells/mL) and incubated for 24 h (or until 95% monolayer confluency was achieved). Monolayers were fixed with 4% paraformaldehyde (PFA) (20 min, room temperature). Once fixed, PFA was removed, and monolayers washed twice with 200 µL sterile 1×PBS. Monolayers were submerged in 1×PBS and stored at 4°C and used within two weeks.

### 2.3. Biofilm Formation

*E. coli* biofilms were formed on the bottom of 96-well microtiter plates with and without fixed keratinocyte monolayers as the substratum. Plate wells were inoculated with 150 µL of diluted overnight bacterial culture (≈1×10^6^ CFU/mL) and incubated for 24 h (37°C, 50 rpm).

### 2.4. Plasma Activated Water Generation and Treatment

Plasma activated water (PAW) was generated as previously described using a bubble spark discharge (BSD) reactor [15] (Fig. 1A). This reactor comprised a stainless-steel metal rod as the high voltage electrode. It is enclosed in a glass sheath with four 0.4 mm diameter holes at the end of the electrode that permit plasm gas to enter the liquid as bubbles. The reactor was placed in 250 mL Schott bottles containing 100 mL of autoclave sterilised Milli-Q water. Using a Leap100 high voltage power supply (PlasmaLeap Technologies, Australia), a voltage input of 150 V, discharge frequency of 1500 Hz, resonance frequency of 60 kHz, and a duty cycle of 100 µs was applied for 20 min with airflow at 1 standard litre per min (slm). As a control, 100 mL autoclave sterilised Milli-Q water was subjected to 20 min exposure to air flow at 1 slm without plasma discharge. Treatment of biofilms grown on both plastic substratum and fixed keratinocyte monolayers used 200 µL of the freshly produced PAW or control to the wells for 15 min (Fig. 1B).

**Figure 1:**
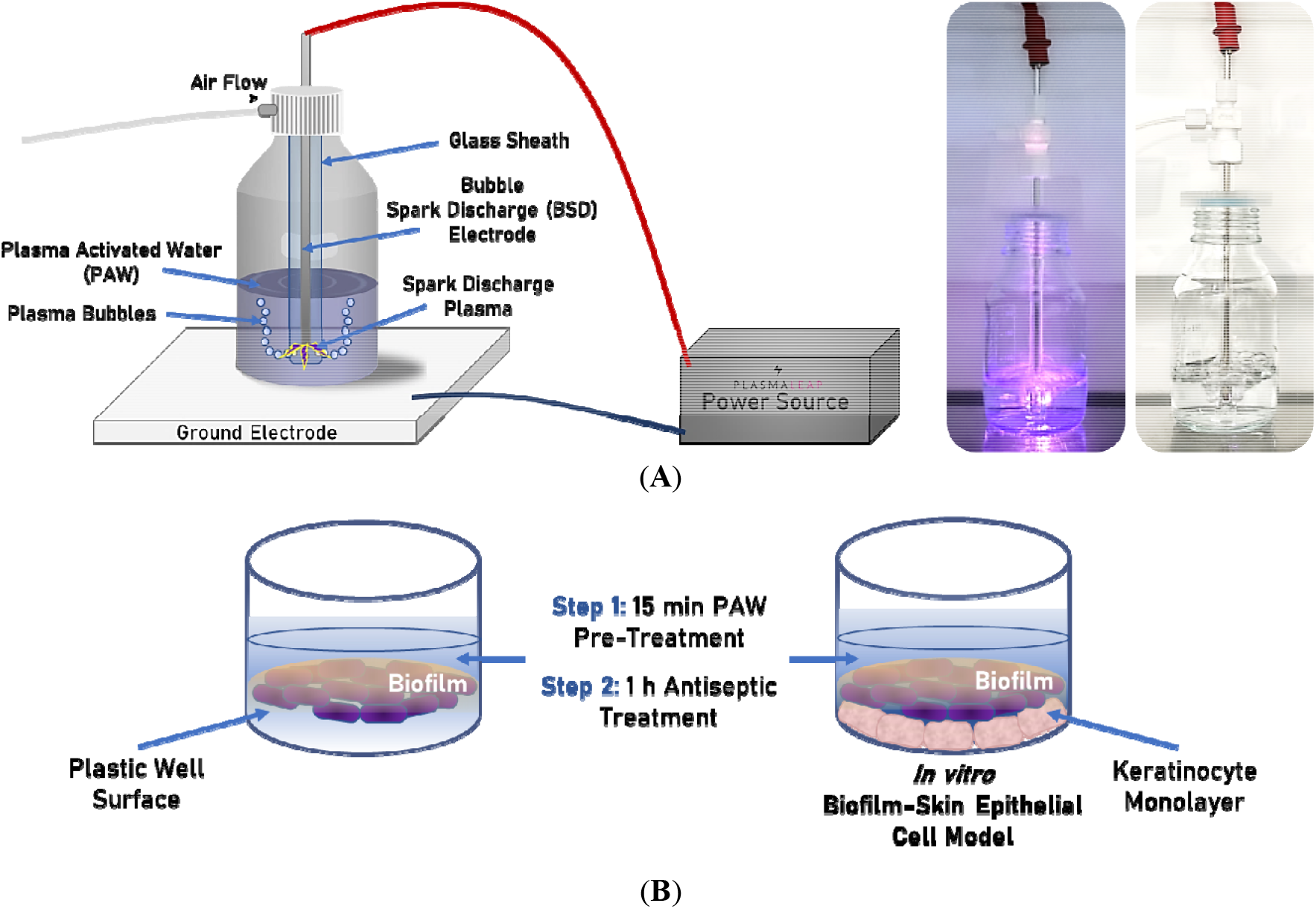
PAW generation and treatment of biofilms. **A)** Schematic representation of the BSD reactor used to generate the PAW with photograph of PAW generation (left) and control generated without plasma discharge (right). **B)** PAW was added directly onto the 24 h *E. coli* biofilms formed on either the plastic well surface (left) or a fixed keratinocyte monolayer (*in vitro* biofilm-skin epithelial cell model; right). PAW was applied for 15 min as a pre-treatment, then biofilms are challenged with clinically relevant topical antiseptics routinely used for treating chronic wounds.

### 2.5. Antimicrobial Agents

Three topical antiseptics routinely used for the treatment of chronic wounds were used: polyhexamethylene biguanide (PHMB) (All Chemical, Australia), povidone iodine (PI) (Sigma-Aldrich, Australia), and a commercially available medical-grade manuka honey (MediHoney, Comvita Ltd, New Zealand). The MediHoney was stored in the dark at 4°C and dissolved in sterile Milli-Q water for use at a stock solution of 40%. Gramicidin (Sigma-Aldrich, Australia) and colistin sodium methanesulfonate (colistin; Sigma-Aldrich, Australia) are antimicrobials with membrane activity and were used as the positive controls for the membrane assays, where appropriate [16].

### 2.6. Antimicrobial Susceptibility Testing

#### 2.6.1. Planktonic Cells - Minimum Inhibitory Concentration and Minimum Bactericidal Concentration Assays

To assess the antimicrobial efficacy of PHMB, PI, MediHoney, PAW, and control against planktonic *E. coli*, minimum inhibitory concentration (MIC) and minimum bactericidal concentration (MBC) assays were performed. The MIC determines the lowest concentration of the antimicrobial that will inhibit visible growth, whilst the MBC is the lowest concentration of an antibacterial agent required to kill *E. coli* cells upon spot platting on LB agar. Standard protocols of either microbroth dilution series (as per CLSI guidelines) or resazurin staining [17, 18] were performed against planktonic suspensions of *E. coli* (≈1×10^6^ CFU/mL), incubated for 24 h at 37°C. Lastly, to determine if these were bactericidal or bacteriostatic against planktonic *E. coli* cells, MBC/MIC ratios were calculated. An MBC/MIC ratio ≤4 was considered bactericidal, whilst an MBC/MIC ratio >4 was considered bacteriostatic [19].

#### 2.6.2. Biofilms - Minimum Biofilm Eradication Concentration Assay

Minimum biofilm eradication concentration (MBEC) assays were utilised to assess *E. coli* biofilm antimicrobial susceptibility. Briefly, the biofilms were washed once with 150 µL Milli-Q water and pre-treated with 200 µL PAW (or control) for 15 min. The PAW (or control) was then removed and the biofilms challenged with 100 µL of two-fold serial dilutions of respective antiseptic (PHMB, PI, or MediHoney) for 1 h, at 37°C. Biofilms were washed, resuspended in sterile 1×PBS, and viable biofilm cells were enumerated via 10-fold serial dilutions and spot plating on LB agar (overnight, 37°C) for subsequent colony counting and CFU/mL determination. The MBEC was determined as the lowest concentration of antimicrobial required to induce complete biofilm eradication, i.e., where 100% of biofilm-associated *E. coli* cells have been killed.

### 2.7. PAW Physicochemical Analysis

The physicochemical properties of PAW and control such as temperature, pH, oxidation-reduction potential (ORP), electrical conductivity, as well as the concentrations of ozone, hydrogen peroxide, nitrite, and nitrate generated via the BSD reactor, were measured as per Rothwell [15] and Zhou [20]. Briefly, a double junction, gel-filled pH probe with built-in temperature sensor was used to measure the pH, ORP was measured using a combination ORP electrode and general-purpose reference electrode, conductivity was measured via a four-ring electrical conductivity probe. These probes and the research-grade benchtop meter were sourced from Hanna Instruments (USA). Dissolved ozone concentrations were determined using a colorimetric assay using the N,N-diethyl-p-phenylenediamine method (accurate at 0.00-2.00 mg/L) with the intensity of the solution at 525 nm measured by a multiparameter benchtop photometer from Hanna Instruments. Hydrogen peroxide was quantified using the titanium (IV) oxysulfate method, measuring the yellow complex formed at 407 nm. Nitrite was quantified using the Griess Reagent method by absorption at 526 nm. Nitrate ions were quantified using a 930 compact Ion Chromatograph (IC) with ProfIC autosampler and automated dilution module (Metrohm). A Metrosep A Supp 7 (5 µm packing, 4 × 250mm) column was used to separate analytes over 32 min using an isocratic flow rate of 0.7 mL/min of 3.6 mmol/L sodium carbonate. Samples were automatically diluted by the instrument 1:20 before injecting 1 μL to the column to ensure peak symmetry.

### 2.8. PAW Physicochemical Impact on Biofilms

Scavengers were used to investigate the effect of specific reactive species to determine which components contribute to the anti-biofilm activity of PAW. The reactive species targets and scavengers that were quenched included superoxide ions using 20 mM disodium 4,5-dihydroxybenzene-1,3-disulfonate (tiron), ozone using 0.1 mM uric acid (can also scavenge hydroxyl radicals) and a general reactive oxygen species (ROS) scavenger (superoxide ions, ozone, hydroxyl radicals) using 20□M ascorbic acid [15]. These scavengers were directly added to the Schott bottles containing 100 mL sterile Milli-Q water prior to PAW generation. A control (no plasma generation) was also included.

As PAW generation is both an acidifying and heat-inducing process, the impact of pH and temperature was also assessed. Biofilms were exposed to sterile Milli-Q water that was adjusted to a pH of 2.8 using nitric acid, and Milli-Q water heated to 51.3°C (the maximum temperature reached during PAW generation), as well as the combination of pH 2.8 Milli-Q water heated to 51.3°C.

### 2.9. Quantification of Biofilm RONS

To further confirm the intracellular accumulation of both ROS and reactive nitrogen species (RNS) upon PAW treatment, biofilms were stained with 20 μM 2’,7’–dichlorofluorescin diacetate (DCFDA; Sigma-Aldrich, Australia) and 5 μmol 4,5-diaminofluorescein diacetate (DAF-FM; Sigma-Aldrich, Australia), respectively [21, 22]. Biofilms were challenged for 15 min with 200 μL PAW and control as above. Once challenged, biofilms were stained with either 150 μL DCFDA or DAF-FM solution for 30 min. The ROS and RNS were detected at an excitation/emission of 485-15 nm/535-15 nm and 495-15 nm/515-15 nm (CLARIOStar), respectively.

### 2.10. Effect of PAW on Membrane Activity

#### 2.10.1. Membrane Depolarisation

Membrane depolarisation was assessed in *E. coli* using 2 µmol/L 3,3′-diethylthiadicarbocyanine iodide (DiSC3(5); Sigma-Aldrich, Australia) [22], a fluorogenic dye measuring changes in transmembrane potential. The dye was allowed to incorporate into 50 µL planktonic *E. coli* cells (≈5×10^6^ CFU/mL) for 20 min at 37°C. Once washed, the cells were exposed to 200 µL PAW and control (0-15 min). As a positive control, 50 µg/mL gramicidin was used. Fluorescence was measured at 600-15/660-15 nm (CLARIOStar), and membrane depolarisation was reported as arbitrary units.

#### 2.10.2. Inner Membrane Permeability

To assess the inner membrane permeability of the *E. coli* cells post-PAW treatment, an ortho-nitrophenyl-β-galactoside (ONPG; Sigma-Aldrich, Australia) assay was performed as per Brun [22]. Planktonic *E. coli* cells were prepared to a final density of ≈5×10^6^ CFU/mL. In a 96-well plate, 50 µL of *E. coli* cells was exposed to 1.5 mM ONPG (dissolved in 1×PBS). Stained *E. coli* cells were then challenged with 200 µL PAW and control. Gramicidin was used as the positive control. ONPG was measured in a time-dependent manner (0-15 in) at 405 nm (CLARIOStar) to determine the inner membrane permeability.

#### 2.10.3. Outer Membrane Permeability

PAW-induced outer membrane permeability was measured based on fluorescent dye N-phenyl-1-naphthylamine (NPN; Sigma-Aldrich, Australia) uptake [22]. 50 µL of planktonic *E. coli* cells (≈5×10^6^ CFU/mL) were mixed with 10 µM NPN and challenged by 200 µL PAW or control. The positive control was 200 µg/mL colistin. NPN-associated fluorescence was measured over time (0-15 min) at excitation/emission wavelengths of 350-15 nm/420-15 nm. At each time point, the value of fluorescence was converted as the percentage of NPN uptake over the observed fluorescence on untreated *E. coli* using *Equation 1* [23]:

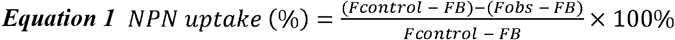

*F*_*obs*_ is the observed fluorescence of NPN with *E. coli* in the presence of PAW or control at a certain time point.

*F*_*control*_ is the fluorescence of NPN with *E. coli* cells in Milli-Q water.

*F*_*B*_ is the background fluorescence in the absence of NPN.

#### 2.10.4. Scanning Electron Microscopy

Scanning electron microscopy (SEM) imaging was utilised to assess the morphological changes induced of *E. coli* biofilm cells following PAW treatment. *E. coli* biofilms were grown for 24 h on 13 mm plastic Nunc Thermanox coverslips (Proscitech, Rochester, USA) in a 12-well polystyrene plate. Biofilms were treated with PAW and control (and positive controls gramicidin and colistin) for 1 and 15 min. Biofilms were air dried and prepared for SEM imaging using methods adapted from Vyas [24] with the following modifications. Biofilms were pre-fixed for 30 min at 4°C, followed by fixation for 1 h at 4°C. Post-fixation, washed biofilms were dehydrated via graded ethanol series (30%, 50%, 70%, 90%, and 3 × 100%) and critical point dried. Dried biofilms were sputter coated with 20 nm platinum (Edwards Vacuum coater, USA) and visualised using a JEOL JSM-7500 microscope (JEOL, Japan) at 500 and 5,000× magnification. Images were taken at random positions to reduce bias.

### 2.11. Statistical Analysis

Statistical analysis was performed using GraphPad Prism (version 8.4.0, GraphPad Software, USA). Experiments were performed in triplicate (with two technical replicates each) and values were expressed as mean ± standard error of the mean (or standard deviation, where appropriate). A one- or two-way ANOVA was performed where appropriate with a Tukey’s multiple comparisons post-hoc test, and P ≤ 0.05 was considered significant.

## 3. Results

### 3.1. PAW Pre-Treatment Greatly Enhances the Anti-Biofilm Activity of Topical Antiseptics

The effectiveness of three topical antiseptics (PHMB, PI, and MediHoney) routinely used to treat chronic wounds was individually assessed against planktonic *E. coli* cells and their MIC, MBC, and MBC/MIC values determined (Table 1). PHMB and PI were both potent bactericidal agents (MBC/MIC≤4), with MIC values of 0.001% and 0.063%, respectively. MediHoney required a much higher dose to inhibit *E. coli* growth (MIC of 10%) and was bacteriostatic (MBC/MIC>4). PAW demonstrated bactericidal activity (MBC/MIC≤4), with a MIC of 3.13%. The Milli-Q water without plasma (termed the control) had no antimicrobial effect (MIC>50%).

**Table 1:** Antimicrobial susceptibility testing of PHMB, PI, MediHoney, PAW, and control against planktonic *E. coli*. MBC/MIC ≤4 is bactericidal, whilst MBC/MIC ratio >4 is bacteriostatic.

| | MIC (%) | MBC (%) | $\frac{MBC}{MIC}$ |
| --- | --- | --- | --- |
| **PHMB** | 0.001 | 0.001 | $\leq 4$ |
| **PI** | 0.063 | 0.25 | $\leq 4$ |
| **MediHoney** | 10 | $>20$ | $>4$ |
| **PAW** | 3.13 | 0.097 | $\leq 4$ |
| **Control** | $>50\%$ | $>50\%$ | $>4$ |

PAW was assessed as a pre-treatment strategy against 24 h *E. coli* biofilms formed on a plastic substratum, followed by treatment with a dilution series of one of the topical antiseptics (Fig. 2). Specifically, PAW was applied first to the biofilms for 15 mins, and then the biofilm further challenged for 1 h with PHMB, PI, or MediHoney. PAW+PHMB and PAW+PI (Fig. 2A and B) completely eradicated biofilm cells at all concentrations tested (MBEC values of PAW+0.001% PHMB and PAW+0.004% PI). These MBEC’s suggest that *E. coli* biofilm susceptibility is either equivalent to, or far exceeds, its planktonic cell counterparts when compared to the PHMB and PI MIC values, respectively. The control treatment (pre-treatment with Milli-Q water without plasma activation) required substantially higher concentrations of PHMB and PI to achieve complete biofilm eradication (MBEC’s control+0.016% PHMB and control+0.063% PI, respectively). For MediHoney, complete biofilm eradication was achieved for PAW pre-treated biofilms (MBEC of PAW+2.5% MediHoney), with biofilm susceptibility far exceeding the planktonic MIC for MediHoney alone (10%). PAW alone was assessed (purple dotted line, Fig 2A-C), consistently reducing biofilm viability by ≈4.5 log when compared to the control (≈7.4 log, blue dotted line, Fig 2A-C).

**Figure 2:**
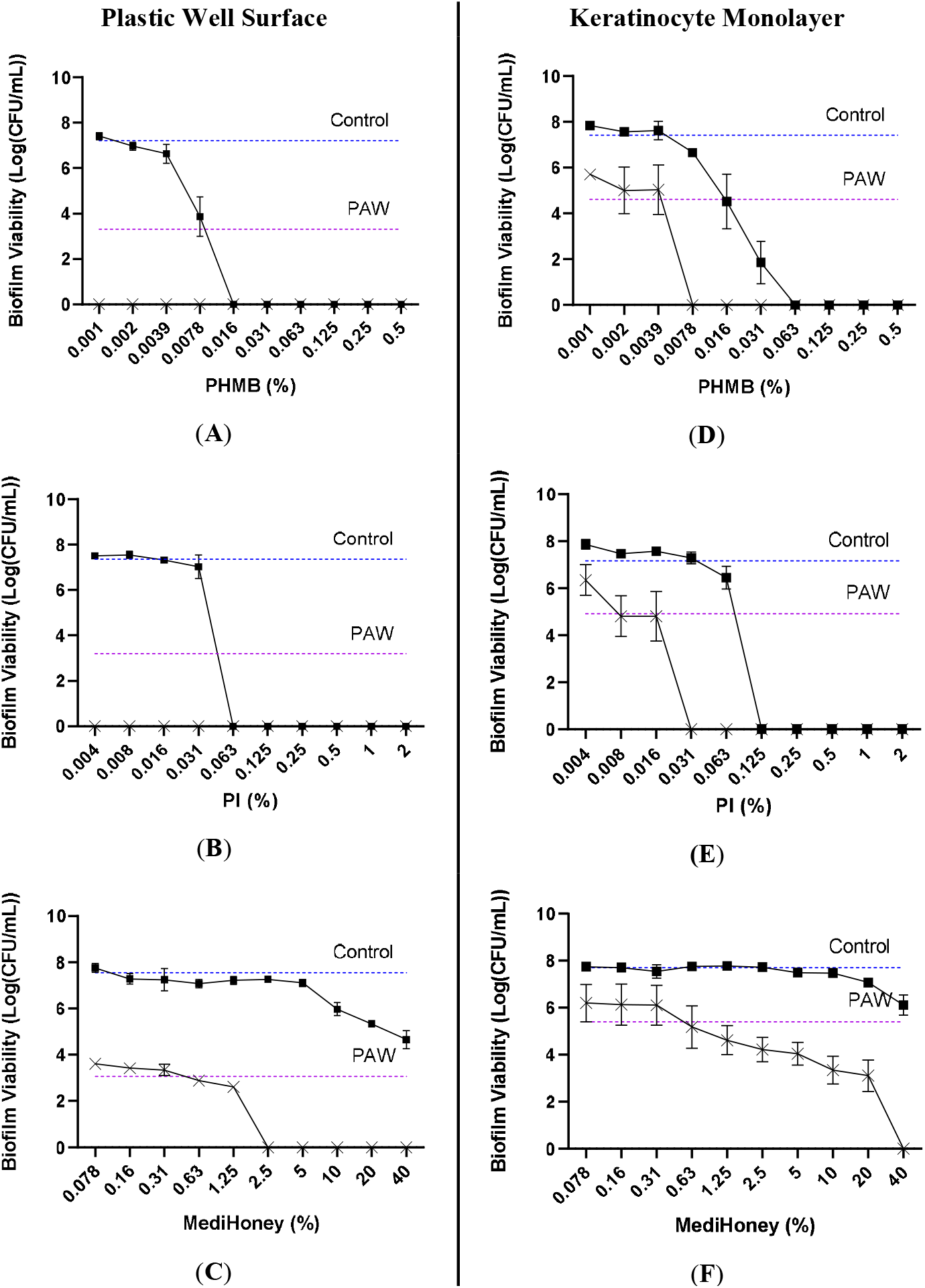
PAW pre-treatment greatly increases the *in vitro* antimicrobial susceptibility of *E. coli* biofilms. Effect on biofilm viability of PAW+PHMB/PI/MediHoney (×), control+PHMB/PI/MediHoney (■), PAW (purple dotted line), and control (blue dotted line) on **A-C**) plastic and **D-F**) keratinocyte monolayer is demonstrated. Data represents mean ± SEM; n = 3 biological replicates, with 2 technical replicates each.

Given the anti-biofilm efficacy of PAW as a pre-treatment on the plastic substratum, this analysis was extended to an *in vitro* biofilm-skin epithelial cell model comprising a keratinocyte monolayer as the substratum for *E. coli* biofilm growth. The efficacy of the antimicrobial treatment was lower for biofilms formed on the keratinocyte monolayer than those formed on plastic (Fig 2D-F). PAW pre-treatment followed by either PHMB or PI (Fig. 2D and E) completely eradicated the biofilm, producing MBECs of PAW+0.0078% PHMB and PAW+0.031% PI, while the control treatment had MBECs of control+0.063% PHMB and control+0.125% PI (Fig 2D and E). PAW+MediHoney (Fig. 2F) achieved complete biofilm eradication at the highest concentration tested (MBEC of PAW+40% MediHoney). Control+MediHoney (Fig. 2F) reduced biofilm viability by ≈1.5 log compared to the control alone (≈7.4 log, blue dotted line, Fig 2D-F). As with the plastic substratum, PAW alone (purple dotted line, Fig 2D-F) did not completely eradicate the biofilms formed on the keratinocyte monolayer but achieved ≈2 log reduction in biofilm viability when compared to the control (blue dotted line).

### 3.2. RONS Primarily Contribute to the Anti-Biofilm Activity of PAW

To determine the mechanisms behind the anti-biofilm activity of PAW, the properties of PAW were investigated, including temperature, pH, oxidation-reduction potential (ORP), conductivity, and RONS (ozone, hydrogen peroxide, nitrite, and nitrate) (Supplemental Table. 1). The PAW was found to have a low pH (pH 2.8) and an initial temperature of 51.3°C, compared to the control (pH 6.2 and 24.2°C). PAW was also notably more conductive (763.3 μS/cm) with a high ORP (502 mV) compared to control (4.8 μS/cm and 390 mV). The RONS that were detected included ozone (1.9 ppm, approaching upper detection limit of 2 ppm), hydrogen peroxide (8.8 ppm), and nitrate (123.0 ppm), while nitrite was not detected (0.0 ppm). RONS were not detected within the control. The effects of pH and temperature were assessed both individually and combined. Neither had significant impacts on *E. coli* biofilm viability (Supplemental Fig. 1). This suggested that the anti-biofilm activity of PAW was primarily due to RONS.

A scavenger assay was performed to determine which reactive species contributed to the anti-biofilm activity of PAW using tiron (superoxide scavenger), uric acid (ozone scavenger), and ascorbic acid (general ROS scavenger). These were added immediately prior to PAW generation and the resulting PAW was then applied to the *E. coli* biofilms for 15 min, with biofilm viability determined via cell enumeration. Scavenging of superoxide, ozone, and ROS generally during the PAW generation process resulted in an increase in *E. coli* biofilm cell viability of ≈1.5 (P ≤ 0.05), 2.5 (P ≤ 0.001), and 7 log (P ≤ 0.0001) respectively, compared to the biofilm viability post-PAW treatment (Fig. 3A).

**Figure 3:**
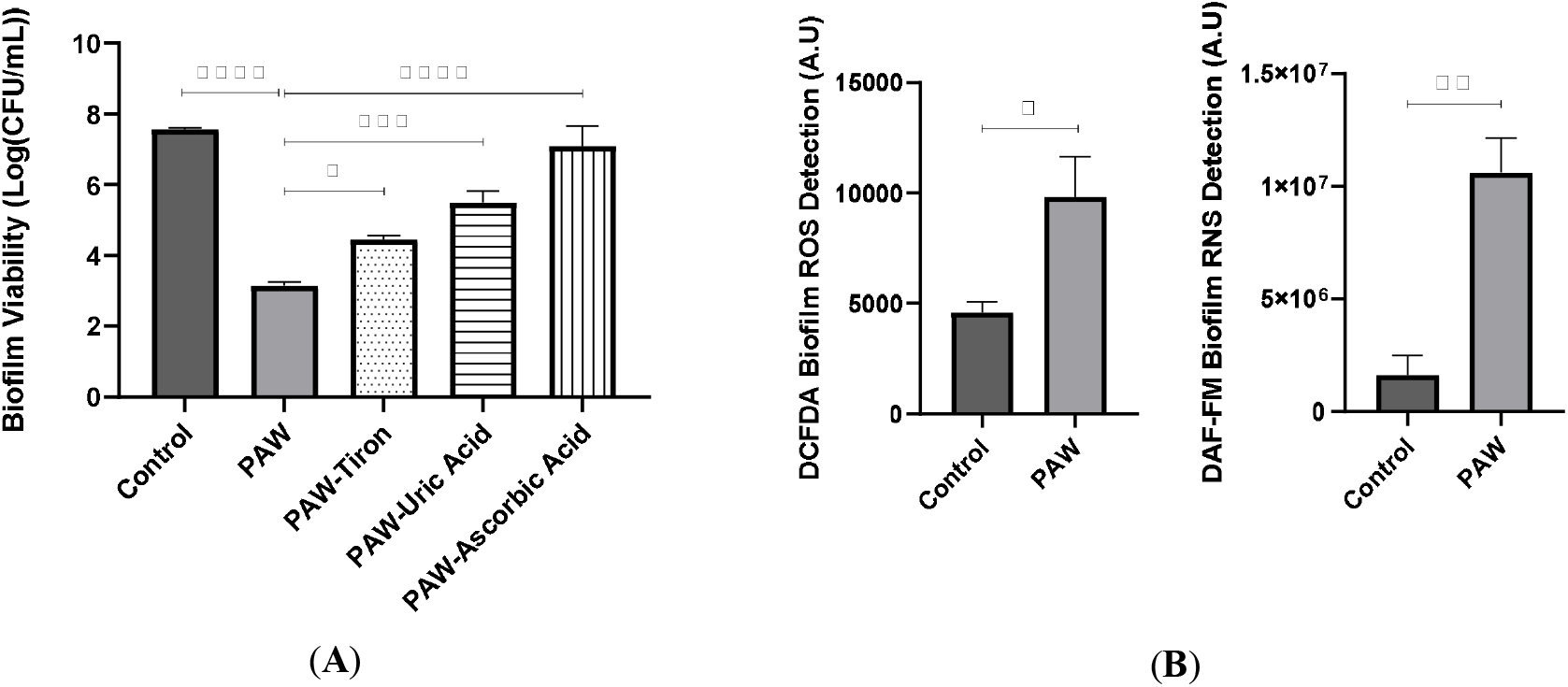
RONS primarily contribute to the anti-biofilm activity of PAW. **A)** PAW with the addition of tiron, uric acid, and ascorbic acid to scavenge superoxide, ozone, and general ROS, respectively. **B)** Intracellular ROS was measured using DCFDA staining (left) and intracellular RNS was measured using DAF-FM staining (right). Data represents mean ± SEM, * (P ≤ 0.05), ** (P ≤ 0.01), *** (P ≤ 0.001), and **** (P ≤ 0.0001); n = 3 biological replicates, with 2 technical replicates each.

The accumulation of RONS within the PAW treated *E. coli* biofilms was then determined using fluorescent staining (Fig. 3B). Compared to the control, a significant increase (P ≤ 0.05) in fluorescent intensity was observed for DCFDA stained biofilms treated with PAW, demonstrating the accumulation of ROS within the biofilm following 15 min PAW treatment (Fig. 3B; left). DAF-FM fluorescence increased even more significantly (P ≤ 0.01), demonstrating a higher abundance of RNS within the PAW treated *E. coli* biofilms (Fig. 3B; right).

### 3.3. PAW Treatment Causes Rapid Outer Membrane Permeability and Membrane Depolarisation

To further determine the mode of action of PAW on *E. coli*, membrane activity was investigated utilising specific stains. For membrane depolarisation, DiSC3(5) was used (Fig. 4A) whilst inner and outer membrane activity used ONPG- and NPN-based assays, respectively (Fig. 4B and C). The greatest effects were seen on the outer membrane (Fig. 4C), where within 1 min of exposure to PAW the outer membrane was significantly perturbed (P ≤ 0.0001) as indicated by NPN uptake. This effect increased until 15 min (P ≤ 0.0001) when compared to the control. The membrane was also significantly depolarised at 1 min of PAW treatment

**Figure 4:**
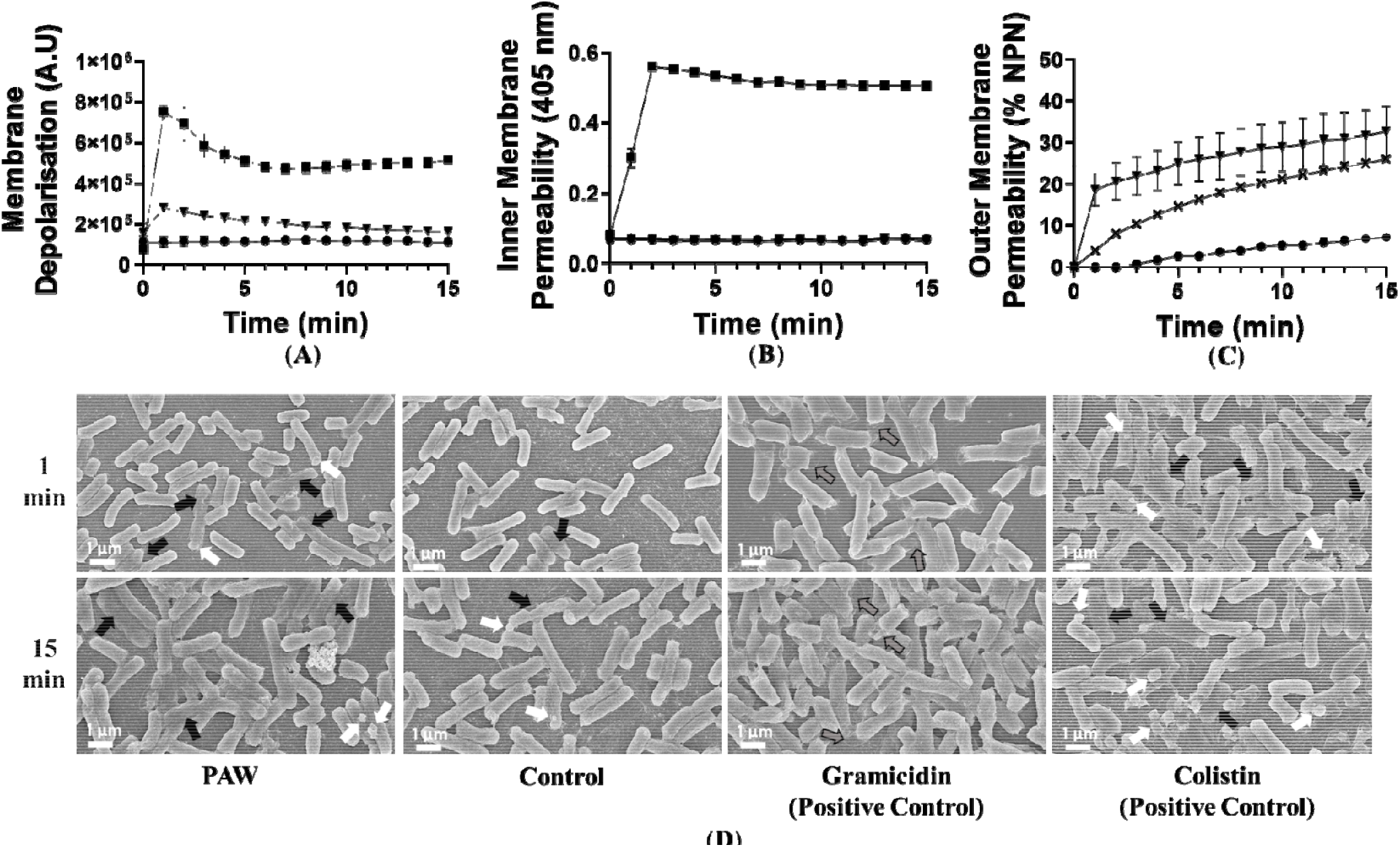
PAW disrupts the integrity of the *E. coli* cell membrane. **A)** Membrane depolarisation was determined for *E. coli* incubated with 2 µm 9 DiSC3(5). *E. coli* cells were challenged with PAW (▾), control (•), and gramicidin (positive control) (■). **B)** Inner membrane permeability was determine 0 the addition of 1.5 mM ONPG to the *E. coli* cells and cytoplasmic b-galactosidase leakage was determined with o-nitrophenol detection (405 nm) challenge with PAW (▾), control (•), and gramicidin (positive control) (■). **C)** Outer membrane permeability was evaluated by incubating *E. coli* with 10 NPN and subsequently challenged with PAW (▾), control (•), and colistin (positive control) (×). NPN uptake was expressed as a percentage (%). **D)** was utilised to visualise morphological changes induced by PAW, particularly on *E. coli* biofilm cell membranes. Biofilms were treated for 1 or 15 min PAW and control. Gramicidin and colistin positive controls were also included. Morphological changes included cell flattening (black arro s), cell memb blebbing (white arrows), and collapsing/concaving inward of individual cell ends (grey arro

(P ≤ 0.0001) (Fig. 4A), but this effect gradually decreased over time and by 11 min depolarisation did not significantly differ from the control. PAW did not appear to induce any inner membrane permeability, as the detected fluorescent values for PAW treated *E. coli* were the same as the control (Fig. 4B).

SEM imaging (Fig. 4D) was conducted to qualitatively distinguish any effects caused by PAW to the *E. coli* cells, particularly in the context of membrane changes. Gramicidin and colistin were also included for comparison. Many of the PAW-treated *E. coli* biofilm cells appeared flattened, with some cells exhibiting membrane blebbing at 1 min of treatment, which was further pronounced at 15 min. Control treated cells (1 and 15 min) also showed flattening, but at a relatively lower frequency (blebbing only seen at 15 min). Both gramicidin and colistin induced extensive morphological changes, and gramicidin was the only treatment to induce significant concaving or collapsing inward of *E. coli* cell ends at 1 min. As with PAW and control treatments, colistin flattened cells and induced prominent cell membrane blebbing at both 1 and 15 min. When inspected at a lower magnification (500 x; Supplemental Fig. 2) the PAW and control treatments did not appear to remove *E. coli* biofilm from the surface, indicating that the PAW generated in this study does not physically dislodge biofilms as part of its mechanism of action.

## 4. Discussion

Biofilm-infected chronic wounds are difficult to treat via conventional antimicrobials [25]. As we fast approach the post-antibiotic era, the development of newer antimicrobials and treatment strategies is critical. Contextually, our results indicate that applying the PAW as an initial wound rinse/soak prior to the topical application of antiseptics (e.g., PHMB, PI, and MediHoney) can aid in the complete eradication of *E. coli* biofilm cells, whilst reducing the concentration of subsequently applied antiseptic. This is important as any remnant surviving biofilm cells can otherwise re-populate and re-establish a biofilm at the wound bed, contributing to recalcitrance and chronicity. Lowering antiseptic concentration can also be beneficial, as some topical antiseptics facilitate dermal hypersensitivity/allergenicity and increase the risk of cytotoxicity for key cell types (e.g., keratinocytes and fibroblasts) which are responsible for wound healing [5]. Further study is needed to investigate the exact synergism occurring between PAW and each antiseptic considering their unique modes of action; PHMB destabilises the microbial membrane; PI disrupts the respiratory chain, disrupts efflux pumps, and denatures cellular proteins and enzymes; and medical-grade honey hinders microbial growth and is rich in antimicrobial ROS (e.g., hydrogen peroxide) [5]. This may provide further insight as to why PAW is more effective when combined with PHMB or PI over MediHoney. Lastly, we demonstrate that PAW pre-treatment is also effective against biofilms generated in the *in vitro* biofilm-skin epithelial cell model that encompasses a keratinocyte monolayer as the substratum for biofilm growth. Several studies have reported that biofilm model choice is crucial when assessing and developing novel antimicrobials and treatment strategies [13]. Biofilms generated in *in vitro* model systems that fail to capture or mimic the infection scenario/local host microenvironment, i.e., in the case of chronic wounds lacking the skin epithelia, three-dimensional tissue layering, or even the wound milieu, may result in biofilms that differ in their architecture/structure, individual biofilm cell morphology, metabolic profile, quorum sensing, as well as their antimicrobial susceptibility (reviewed [13]). Hence, the findings of this study indicates that a more realistic prediction for translatable antimicrobial success under clinical settings is greatly increased and/or achievable.

Considering PAWs demonstrated antimicrobial potency and anti-biofilm efficacy as a pre-treatment strategy, the mechanisms underpinning its activity were investigated. Physicochemical analysis revealed several RONS present within the PAW including ozone, hydrogen peroxide, and nitrate. These reactive species have been found to inactivate several pathogens, some of which have been implicated in chronic wounds (e.g., *E. coli* and *P. aeruginosa*) [26, 27]. Given the abundance and diversity of ROS in PAW, along with their widely recognised role in CAP-mediated microbial damage [28], these were the focus of subsequent study. Firstly, a scavenger assay was performed to selectively remove ROS species. The greatest increase in biofilm viability was seen for PAW scavenged via ascorbic acid, whereby several important ROS (e.g., superoxide, ozone, and several ozone by-products, like hydroxyl radicals) were removed. In fact, scavenging these various ROS from PAW was so effective that *E. coli* biofilm viability did not significantly differ to the biofilm control. Xia [21] found PAW-associated superoxide was crucial for *E. coli* biofilm removal, and Rothwell [15] found superoxide (and/or its downstream reactive species) were primary contributors to PAW-mediated inactivation of planktonic *E. coli* and *Listeria innocua* cells. Saijai [29] found that ozonated bubble water was a strong sterilising agent against *E. coli*. Moreover, ozone can generate several other reactive downstream ROS (e.g., hydroxyl radicals). Hydroxyl radicals are potent antibacterial agents against several planktonic and biofilm bacteria like *E. coli* and *Streptococcus mutans* [29-31]. Taken together, it is apparent that scavenging superoxide and ozone from PAW subsequently prohibits the formation of various ROS by-products. Collectively, their removal significantly reduces the antimicrobial power of PAW.

CAP has previously been shown to inactivate bacterial cells by creating an intracellularly high oxidative stress environment with cells responding to this environment by producing additional RONS [28]. Oxidative stress is harmful to microbial cells and their intracellular components (e.g., nucleic acids, proteins, lipids), and inducing such a surge in intracellular ROS causes irreversible damage and enhances lethality [28, 32, 33]. Patange [28] described several ROS (superoxide, peroxide, hydroxyl radicals) as key proponents in CAP-mediated intracellular damage of *Listeria monocytogenes* biofilm cells. Similarly, PAW-associated hydrogen peroxide, superoxide, ozone, and their by-product ROS may each contribute to a damaging oxidative stress response in the treated *E. coli* biofilms. This may result in an increased intracellular ROS production which is damaging to the cells. PAW-induced oxidative stress can also generate high concentrations of intracellular RNS within Gram-positive and Gram-negative bacterial cells (e.g., *S. aureus, L. monocytogenes*, and *E. coli*) [28, 34]. Here, we also demonstrated significant intracellular RNS accumulation post-PAW treatment, with relatively higher RNS detected than ROS. Additionally, it also possible that PAW-derived RONS directly penetrated through the EPS and accumulated within the biofilm structure [27]. Once in the biofilm structure, PAW-associated RONS can infiltrate into *E. coli* cells by active transport through the lipid bilayer, or more passively through membrane pores [35].

Lastly, the membrane activity of PAW was investigated. Ozone was a prominent potent ROS in our PAW with significant anti-biofilm activity. Komanapalli and Lau [36] observed that short-term ozone exposure (1-5 min), resulted in rapid *E. coli* cellular membrane lipid oxidation and cytoplasmic release of proteins and nucleic acid. Leakage was linked to increased membrane permeability [36]. Ozone-induced membrane lipid oxidation can also cause notable changes to the physical properties of the microbial membrane, e.g., inducing membrane depolarisation [37]. Here, within 1-minute of PAW treatment, *E. coli* cells had significant membrane depolarisation and outer membrane permeability. Hence, ozone may play an important role, thwarting the microbial membrane. Zhang [38] suggests that CAP-induced membrane damage involves the cumulative impact of several long- and short-lived ROS like hydroxyl radicals, hydrogen peroxide, and ozone. These can act on membrane-associated proteins, further triggering oxidative stress within *E. coli* cells, a process resulting in rapid death [38]. SEM imaging of PAW-treated *E. coli* biofilm cells revealed significant morphological changes with cells flattening and membrane blebbing. *In vivo*, several Gram-negative pathogens (e.g., *P. aeruginosa* and *Helicobacter pylori*) have been found to produce outer membrane vesicles (OMVs) that are released as a survival mechanism in response to immune cells like macrophages undergoing “oxidative burst”, where potent antimicrobial ROS is released [39]. OMVs are spherical, extracellular vesicles that bud off from the outer membrane, and when observed under the microscope appear as “blebs” on the microbial surface [39, 40]. *E. coli* has also been shown to produce OMV’s in response to hydrogen peroxide, other ROS, as well as other stressors like increased temperature and hyperosmotic stress [40]. Hence, it is possible that *E. coli* biofilm cell membrane blebbing resulted from both PAW-associated ROS and the other physicochemical properties of the PAW.

## 5. Conclusions

This study highlights the utility of PAW as a pre-treatment strategy, potentiating the efficacy of topical antiseptics that are routinely used in the treatment of infected chronic wounds. Initially, the PAW is likely killing a significant portion of biofilm cells, enhancing the anti-biofilm activity of subsequently applied antiseptics. Importantly, complete eradication is also achievable when biofilms are generated under conditions that encompass host factors, i.e., when grown on keratinocyte monolayers of the *in vitro* biofilm-skin epithelial cell model. Mechanistically, PAW-associated reactive species are pivotal to inducing *E. coli* biofilm cell death, leading to intracellular RONS accumulation and rapid cell membrane abrogation. Overall, this study provides a solid basis for additional investigation into PAW as a pre-treatment strategy for chronic wounds infected by other relevant microbes (e.g., *S. aureus, P. aeruginosa*, and *C. albicans*), and with differing antimicrobials (e.g., topical disinfectants) or treatment strategies (e.g., debridement). PAW is a promising alternative antimicrobial considering the AMR crisis, providing innovation towards effective wound treatment and clinical practice.

## 6. Acknowledgements

This work was supported by the Australian Research Council Discovery Scheme (DP210101358). We thank Prof Martina Sanderson-Smith (University of Wollongong) for kindly providing the HaCaT cell line utilised in this study, and for her overall support of this project. The authors acknowledge the valuable technical assistance and support provided by staff, namely, Dr Mitchell J. B. Nancarrow and Dr Qiang Zhu, at the University of Wollongong Electron Microscopy Centre.

## 7. Author contributions

Conceptualisation, H.K.N.V.; methodology, H.K.N.V.; formal analysis, H.K.N.V.; experimental investigation, H.K.N.V., B.X., D.A., N.P.G., J.G.R.; data curation, H.K.N.V.; visualisation, H.K.N.V.; writing—original draft preparation, H.K.N.V.; writing—review and editing H.K.N.V., B.X., D.A., N.P.G., J.G.R., S.A.R., D.C., P.J.C., A.M-P.; supervision, A.M-P. and H.K.N.V.; resources, A.M-P. and P.J.C.; funding acquisition, A.M-P. All authors have read and agreed to the published version of the manuscript.

## 8. Conflict of Interest

Patrick J. Cullen is the CEO of PlasmaLeap Technologies, the supplier of the plasma power source and BSD reactor utilised in this study.

## Supplementary Data

**Supplementary Table 1:** The physicochemical properties of PAW and control generated for 20 min using the BSD reactor. Data represents mean ± Std Dev, n = 3 replicates.

|  | PAW | control |
| --- | --- | --- |
| Temperature ($^{\circ}$C) | 51.3 ± 1.2 | 24.2 ± 0.4 |
| pH | 2.8 ± 0.0 | 6.2 ± 0.2 |
| ORP (mV) | 502.0 ± 6.1 | 390 ± 14.1 |
| Conductivity (μS/cm) | 763.3 ± 35.1 | 4.8 ± 2.0 |
| Ozone (ppm) | 1.9 ± 0.2 | 0.0 ± 0.0 |
| Hydrogen Peroxide (ppm) | 8.8 ± 1.5 | 0.0 ± 0.0 |
| Nitrite (ppm) | 0.0 ± 0.0 | 0.0 ± 0.0 |
| Nitrate (ppm) | 123.0 ± 4.0 | 0.5 ± 0.1* |

\* Trace quantities detected, likely as a contaminant

**Supplementary Figure 1:**
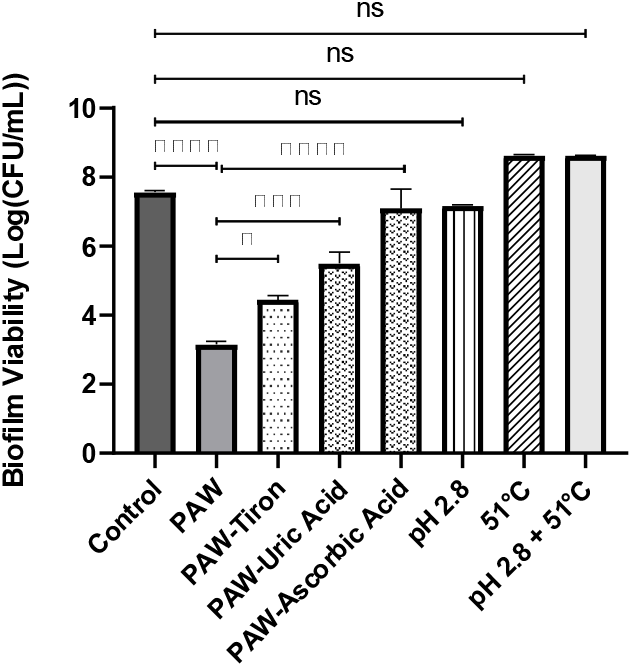
RONS primarily contribute to the anti-biofilm activity of PAW. Addition of tiron, uric acid, and ascorbic acid effectively scavenge superoxide anions, ozone, and general ROS from the PAW, significantly increasing biofilm viability (compared to biofilms treated with whole PAW). Whilst Milli-Q water at pH 2.8, 51°C, and combined pH 2.8 +51°C do not significantly impact biofilm viability, instead closely resemble viability of control.

**Supplementary Figure 2:**
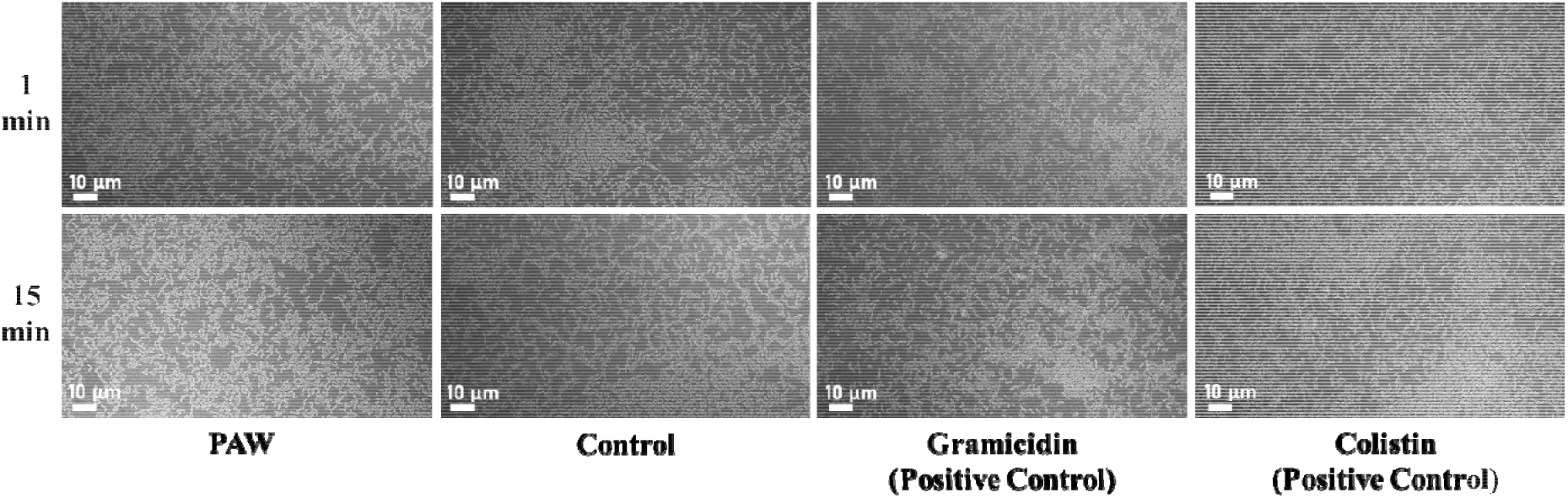
SEM imaging of *E. coli* biofilms treated for 1 and 15 mins with PAW, control, and Positive controls (Gramicidin and Colistin) at 500 x magnification. SEM images demonstrate that neither PAW nor any of the controls, physically dislodge the biofilms. Moreover, SEM images show that biofilm cell numbers/samples are relatively consistent between treatments.

## Notes

### Competing Interest Statement

Co-author, Patrick J. Cullen, is the CEO of PlasmaLeap Technologies, the supplier of the plasma power source and BSD reactor utilised in this study.

## References

[1] Guest JF, Fuller GW, Vowden P. Cohort study evaluating the burden of wounds to the UK’s National Health Service in 2017/2018: update from 2012/2013. BMJ Open. 2020;10:e045253.

[2] McCosker L, Tulleners R, Cheng Q, Rohmer S, Pacella T, Graves N, et al. Chronic wounds in Australia: A systematic review of key epidemiological and clinical parameters. 2019;16:84–95.

[3] Rahim K, Saleha S, Zhu X, Huo L, Basit A, Franco OL. Bacterial Contribution in Chronicity of Wounds. Microbial Ecology. 2017;73:710–21.

[4] Bjarnsholt T, Kirketerp-Møller K, Jensen P∅ Madsen KG, Phipps R, Krogfelt K, et al. Why chronic wounds will not heal: a novel hypothesis. Wound Repair and Regeneration. 2008;16:2–10.

[5] Sibbald RG, Elliott JA, Verma L, Brandon A, Persaud R, Ayello EA. Update: Topical Antimicrobial Agents for Chronic Wounds. Adv Skin Wound Care. 2017;30:438–50.

[6] Puligundla P, Mok C. Potential applications of nonthermal plasmas against biofilm[associated micro[organisms in vitro. Journal of applied microbiology. 2017;122:1134–48.

[7] Gilmore BF, Flynn PB, O’Brien S, Hickok N, Freeman T, Bourke P. Cold plasmas for biofilm control: opportunities and challenges. Trends in biotechnology. 2018;36:627–38.

[8] Weltmann K, Von Woedtke T. Plasma medicine—current state of research and medical application. Plasma Physics and Controlled Fusion. 2016;59:014031.

[9] Mai-Prochnow A, Alam D, Zhou R, Zhang T, Ostrikov K, Cullen PJ. Microbial decontamination of chicken using atmospheric plasma bubbles. 2021;18:2000052.

[10] Nicol MJ, Brubaker TR, Honish BJ, Simmons AN, Kazemi A, Geissel MA, et al. Antibacterial effects of low-temperature plasma generated by atmospheric-pressure plasma jet are mediated by reactive oxygen species. Scientific reports. 2020;10:1–11.

[11] Mai-Prochnow A, Clauson M, Hong J, Murphy AB. Gram positive and Gram negative bacteria differ in their sensitivity to cold plasma. Scientific Reports. 2016;6:38610.

[12] Aboubakr HA, Parra FS, Collins J, Bruggeman P, Goyal SM. Ìn situ inactivation of human norovirus GII. 4 by cold plasma: Ethidium monoazide (EMA)-coupled RT-qPCR underestimates virus reduction and fecal material suppresses inactivation. Food microbiology. 2020;85:103307.

[13] Vyas HKN, Xia B, Mai-Prochnow A. Clinically relevant in vitro biofilm models: A need to mimic and recapitulate the host environment. Biofilm. 2022;4:100069.

[14] Vyas HKN, McArthur JD, Sanderson-Smith ML. An optimised GAS-pharyngeal cell biofilm model. Scientific Reports. 2021;11:8200.

[15] Rothwell JG, Alam D, Carter DA, Soltani B, McConchie R, Zhou R, et al. The antimicrobial efficacy of plasma-activated water against Listeria and E. coli is modulated by reactor design and water composition. 2022;132:2490–500.

[16] Prenner EJ, Lewis RNAH, McElhaney RN. The interaction of the antimicrobial peptide gramicidin S with lipid bilayer model and biological membranes. Biochimica et Biophysica Acta (BBA) - Biomembranes. 1999;1462:201–21.

[17] Humphries R, Bobenchik AM, Hindler JA, Schuetz AN. Overview of Changes to the Clinical and Laboratory Standards Institute Performance Standards for Antimicrobial Susceptibility Testing, M100, 31st Edition. Journal of Clinical Microbiology. 2021;59:10.1128/jcm.00213-21.

[18] Elshikh M, Ahmed S, Funston S, Dunlop P, McGaw M, Marchant R, et al. Resazurin-based 96-well plate microdilution method for the determination of minimum inhibitory concentration of biosurfactants. Biotechnology Letters. 2016;38:1015–9.

[19] Pankey G, Sabath L. Clinical relevance of bacteriostatic versus bactericidal mechanisms of action in the treatment of Gram-positive bacterial infections. Clinical infectious diseases. 2004;38:864–70.

[20] Zhou R, Zhou R, Zhang X, Zhuang J, Yang S, Bazaka K, et al. Effects of Atmospheric-Pressure N2, He, Air, and O2 Microplasmas on Mung Bean Seed Germination and Seedling Growth. Scientific Reports. 2016;6:32603.

[21] Xia B, Vyas HKN, Zhou R, Zhang T, Hong J, Rothwell JG, et al. The importance of superoxide anion for Escherichia coli biofilm removal using plasma-activated water. Journal of Environmental Chemical Engineering. 2023:109977.

[22] Brun P, Bernabè G, Marchiori C, Scarpa M, Zuin M, Cavazzana R, et al. Antibacterial efficacy and mechanisms of action of low power atmospheric pressure cold plasma: membrane permeability, biofilm penetration and antimicrobial sensitization. Journal of applied microbiology. 2018;125:398–408.

[23] Lee DL, Powers JPS, Pflegerl K, Vasil ML, Hancock REW, Hodges RS. Effects of single d-amino acid substitutions on disruption of β-sheet structure and hydrophobicity in cyclic 14-residue antimicrobial peptide analogs related to gramicidin S. The Journal of Peptide Research. 2004;63:69–84.

[24] Vyas HKN, Indraratna AD, Everest-Dass A, Packer NH, De Oliveira DMP, Ranson M, et al. Assessing the Role of Pharyngeal Cell Surface Glycans in Group A Streptococcus Biofilm Formation. 2020;9:775.

[25] Kim PJ, Steinberg JS. Wound Care: Biofilm and Its Impact on the Latest Treatment Modalities for Ulcerations of the Diabetic Foot. Seminars in Vascular Surgery. 2012;25:70–4.

[26] Joshi SG, Cooper M, Yost A, Paff M, Ercan UK, Fridman G, et al. Nonthermal dielectric-barrier discharge plasma-induced inactivation involves oxidative DNA damage and membrane lipid peroxidation in Escherichia coli. Antimicrobial agents and chemotherapy. 2011;55:1053–62.

[27] Alkawareek MY, Algwari QT, Laverty G, Gorman SP, Graham WG, O’Connell D, et al. Eradication of Pseudomonas aeruginosa biofilms by atmospheric pressure non-thermal plasma. PLoS One. 2012;7:e44289.

[28] Patange A, O’Byrne C, Boehm D, Cullen PJ, Keener K, Bourke P. The Effect of Atmospheric Cold Plasma on Bacterial Stress Responses and Virulence Using Listeria monocytogenes Knockout Mutants. Front Microbiol. 2019;10:2841.

[29] Saijai S, Thonglek V, Yoshikawa K. Sterilization effects of ozone fine (micro/nano) bubble water. Int J Plasma Environ Sci Technol. 2019;12:55–8.

[30] Nakamura K, Shirato M, Kanno T, Örtengren U, Lingström P, Niwano Y. Antimicrobial activity of hydroxyl radicals generated by hydrogen peroxide photolysis against Streptococcus mutans biofilm. International Journal of Antimicrobial Agents. 2016;48:373–80.

[31] Wang H, Hasani M, Wu F, Prosser R, MacHado GB, Warriner K. Hydroxyl-radical activated water for inactivation of Escherichia coli O157:H7, Salmonella and Listeria monocytogenes on germinating mung beans. International Journal of Food Microbiology. 2022;367:109587.

[32] Wang X, Zhao X. Contribution of Oxidative Damage to Antimicrobial Lethality. Antimicrobial Agents and Chemotherapy. 2009;53:1395–402.

[33] Kohanski MA, Dwyer DJ, Hayete B, Lawrence CA, Collins JJ. A common mechanism of cellular death induced by bactericidal antibiotics. Cell. 2007;130:797–810.

[34] Borkar SB, Negi M, Kaushik N, Abdul Munnaf S, Nguyen LN, Choi EH, et al. Plasma-Generated Nitric Oxide Water Mediates Environmentally Transmitted Pathogenic Bacterial Inactivation via Intracellular Nitrosative Stress. Int J Mol Sci. 2023;24.

[35] Han L, Patil S, Boehm D, Milosavljević V, Cullen PJ, Bourke P. Mechanisms of Inactivation by High-Voltage Atmospheric Cold Plasma Differ for Escherichia coli and Staphylococcus aureus. Applied and Environmental Microbiology. 2016;82:450–8.

[36] Komanapalli IR, Lau BHS. Ozone-induced damage of Escherichia coli K-12. Applied Microbiology and Biotechnology. 1996;46:610–4.

[37] Brodowska AJ, Nowak A, Śmigielski K. Ozone in the food industry: Principles of ozone treatment, mechanisms of action, and applications: An overview. Critical reviews in food science and nutrition. 2018;58:2176–201.

[38] Zhang H, Ma J, Shen J, Lan Y, Ding L, Qian S, et al. Roles of membrane protein damage and intracellular protein damage in death of bacteria induced by atmospheric-pressure air discharge plasmas. RSC Adv. 2018;8:21139–49.

[39] Sabra W, Lunsdorf H, Zeng A-P. Alterations in the formation of lipopolysaccharide and membrane vesicles on the surface of Pseudomonas aeruginosa PAO1 under oxygen stress conditions. Microbiology. 2003;149:2789–95.

[40] Schwechheimer C, Kuehn MJ. Outer-membrane vesicles from Gram-negative bacteria: biogenesis and functions. Nature Reviews Microbiology. 2015;13:605–19.

