## Supplemental Table 1 for "Plasma Activated Water as a Pre-Treatment Strategy in the Context of Biofilm-Infected Chronic Wounds"

### **Supplementary Data**

**Supplemental Table 1: The physicochemical properties of PAW and control generated for 20 min using the BSD reactor.** Data represents mean ± Std Dev, n = 3 replicates.

|  | **PAW** | **control** |
| --- | --- | --- |
| **Temperature (°C)** | 51.3 ± 1.2 | 24.2 ± 0.4 |
| **pH** | 2.8 ± 0.0 | 6.2 ± 0.2 |
| **ORP (mV)** | 502.0 ± 6.1 | 390 ± 14.1 |
| **Conductivity (μS/cm)** | 763.3 ± 35.1 | 4.8 ± 2.0 |
| **Ozone (ppm)** | 1.9 ± 0.2 | 0.0 ± 0.0 |
| **Hydrogen Peroxide (ppm)** | 8.8 ± 1.5 | 0.0 ± 0.0 |
| **Nitrite (ppm)** | 0.0 ± 0.0 | 0.0 ± 0.0 |
| **Nitrate (ppm)** | 123.0 ± 4.0 | 0.5 ± 0.1* |

* Trace quantities detected, likely as a contaminant
