## Supplemental Figure 1 for "Plasma Activated Water as a Pre-Treatment Strategy in the Context of Biofilm-Infected Chronic Wounds"

### **Supplementary Data**

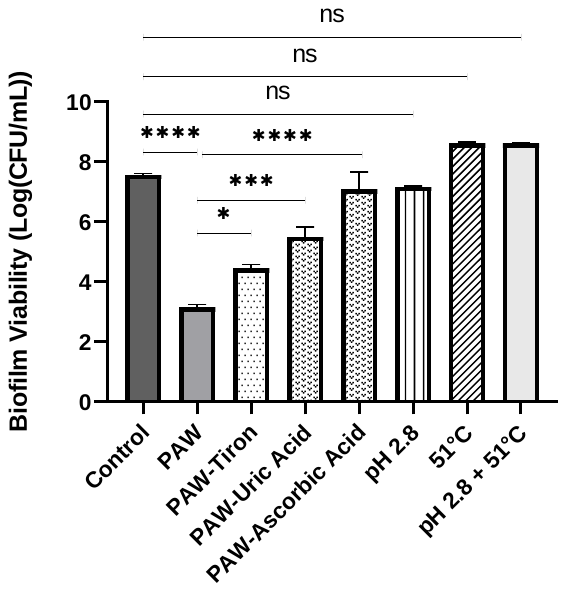

**Supplemental Figure 1: RONS primarily contribute to the anti-biofilm activity of PAW.** Addition of tiron, uric acid, and ascorbic acid effectively scavenge superoxide anions, ozone, and general ROS from the PAW, significantly increasing biofilm viability (compared to biofilms treated with whole PAW). Whilst Milli-Q water at pH 2.8, 51°C, and combined pH 2.8 +51°C do not significantly impact biofilm viability, instead closely resemble viability of control.
