## Supplemental Figure 2 for "Plasma Activated Water as a Pre-Treatment Strategy in the Context of Biofilm-Infected Chronic Wounds"

### **Supplementary Data**

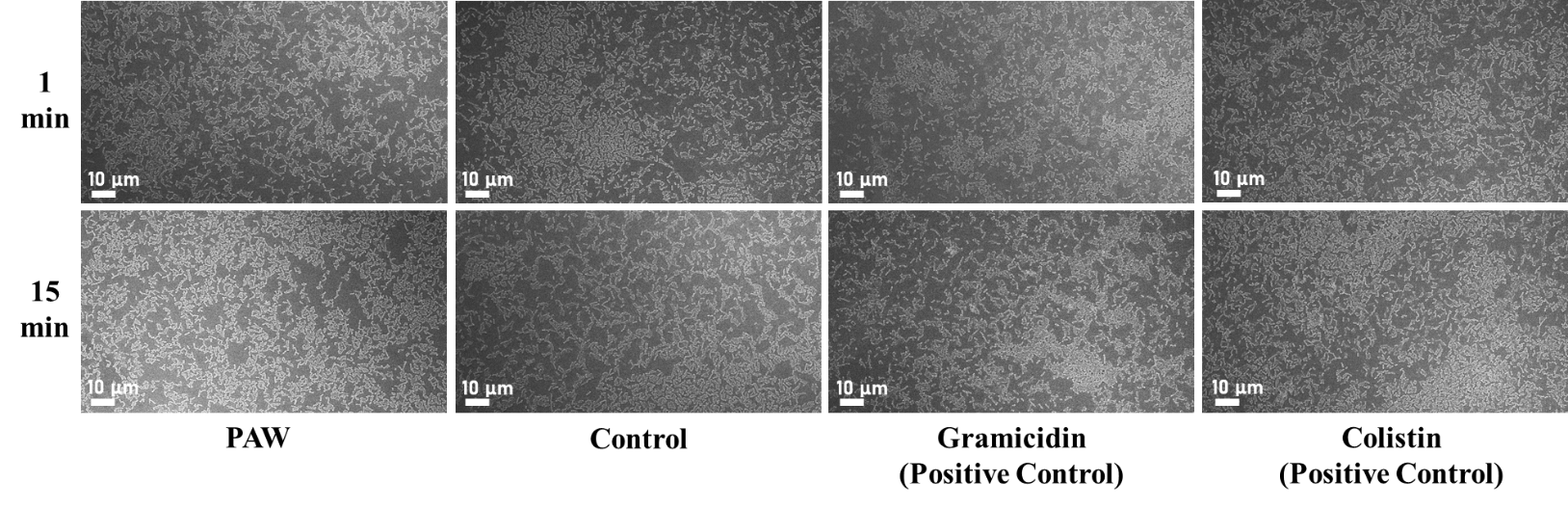

**Supplemental Figure 2: SEM imaging of *E. coli* biofilms treated for 1 and 15 mins with PAW, control, and Positive controls (Gramicidin and Colistin) at 500 x magnification.** SEM images demonstrate that neither PAW nor any of the controls, physically dislodge the biofilms. Moreover, SEM images show that biofilm cell numbers/samples are relatively consistent between treatments.
